# Rescue of the first Alphanucleorhabdovirus entirely from cloned complementary DNA: an efficient vector for systemic expression of foreign genes in maize and insect vectors

**DOI:** 10.1101/2022.05.28.493294

**Authors:** Surapathrudu Kanakala, Cesar Augusto Diniz Xavier, Kathleen M. Martin, Hong Hanh Tran, Margaret G. Redinbaugh, Anna E. Whitfield

**Author notes:** **Correspondence** Anna E. Whitfield, Department of Entomology and Plant Pathology, North Carolina State University, Raleigh, North Carolina 27606. Kathleen M. Martin, Department of Entomology and Plant Pathology, Auburn University, AL, USA 36849. Hong Hanh Tran, Department of Pathology and Laboratory Medicine, Schulich School of Medicine & Dentistry, Western University, London, ON, Canada N6A 3K7.

## Abstract

Recent reverse genetics technologies have enabled genetic manipulation of plant negative-strand RNA virus (NSR) genomes. Here, we report construction of an infectious clone for the maize-infecting *Alphanucleorhabdovirus maydis*, the first efficient NSR vector for maize. The full-length infectious clone was established using agrobacterium-mediated delivery of full-length maize mosaic virus (MMV) antigenomic RNA and the viral core proteins (nucleoprotein N, phosphoprotein P, and RNA-directed RNA polymerase L) required for viral transcription and replication into *Nicotiana benthamiana*. Insertion of intron 2 *ST-LS1* into the viral L gene increased stability of the infectious clone in *Escherichia coli* and *Agrobacterium tumefaciens*. To monitor virus infection *in vivo*, a GFP gene was inserted in between the N and P gene junctions to generate recombinant MMV-GFP. cDNA clones of MMV-WT and MMV-GFP replicated in single cells of agroinfiltrated *N. benthamiana*. Uniform systemic infection and high GFP expression were observed in maize inoculated with extracts of the infiltrated *N. benthamiana* leaves. Insect vectors supported virus infection when inoculated via feeding on infected maize or microinjection. Both MMV-WT and MMV-GFP were efficiently transmitted to maize by planthopper vectors. The GFP reporter gene was stable in the virus genome and expression remained high over three cycles of transmission in plants and insects. The MMV infectious clone will be a versatile tool for expression of proteins of interest in maize and cross-kingdom studies of virus replication in plant and insect hosts.

## 1. INTRODUCTION

Viruses can be used as multifaceted recombinant vectors for virus-induced gene silencing (VIGS), heterologous protein expression and genome editing in both dicotyledonous and monocotyledonous plants. However, efficient, stable, and systemic expression of heterologous proteins at high levels by many virus-based vectors in their hosts is challenging due to genetic instability (reviewed in Willemsen & Zwart, 2019). The *Rhadoviridae* is a diverse family of negative-strand RNA viruses (NSRs) that infect vertebrates, invertebrates, and plants. Plant non-segmented rhabdoviruses are currently classified into four genera, *Cytorhabdovirus* that replicates in the cytoplasm and three nuclear replicating genera *Alphanucleorhabdovirus, Betanucleorhabdovirus*, and *Gammanucleorhabdovirus* (Dietzgen et al., 2020). The genome of plant rhabdoviruses consists of 3’ leader and 5’ trailer sequences and five viral structural protein genes organised in the order of [3’-nucleoprotein (N), phosphoprotein (P), matrix protein (M), glycoprotein (G), and viral polymerase (L)-5’] and separated by conserved gene junctions (Jackson & Li, 2016; Walker et al., 2011). The genomic RNA (gRNA) is complementary to its mRNA, however it requires a minimal infectious unit consisting of gRNA, N, P and L proteins for initiation of virus transcription and replication. Unlike animal rhabdoviruses, plant-adapted rhabdoviruses have at least one additional protein that mediates cell-to-cell movement in plant hosts (Jackson & Li, 2016).

Constructing NSR virus infectious clones continues to be a technical challenge due to large genome sizes, ranging from 10-16 kb in size, and NSR genome is not infectious without the nucleocapsid core, which consists of N, P, and L proteins; thus, the full-length genome and these proteins must be delivered into the same cell. Nevertheless, rhabdoviruses have several features that make them excellent viral vectors: i) virions are bacilliform and can encapsidate an expanded genome, ii) polar transcriptional mechanism of mRNAs provides regulated expression of cargos in the viral genome, and iii) the inserts are stable due to their low rate of genome recombination (Finke and Conzelmann, 2005; Jackson & Li, 2016; Roberts & Rose, 1999). The first infectious clones for NSRs were developed for the animal rhabdoviruses, Rabies virus (Schnell et al., 1994) and Vesicular stomatitis virus (VSV) (Lawson et al., 1995). Cells were cotransfected with antigenomic RNA (agRNA) and viral replication proteins (N, P and L) with transcription facilitated by bacteriophage T7 polymerase.

In the case of plant NSRs, the path for generating infectious virus from cDNA clones required overcoming additional difficulties. The lack of continuous cell cultures for plant NSRs made this technology more challenging, and the relative rigidity of the plant cell wall hinders the entry of multiple plasmids into a single cell to express the full-length genome and replication proteins (N, P and L) required for launching plant NSRs (Walpita & Flick 2005). Additionally, all plant NSR infectious clones to date require co-expressing viral suppressors of RNA silencing (VSRs: tomato bushy stunt virus (TBSV) p19, tobacco etch virus (TEV) HcPro and barley stripe mosaic virus (BSMV) γb proteins) that interfere with host RNA silencing to enhance ribonucleoproteins (RNPs) in *trans* (Ganesan et al., 2013; reviewed in German et al., 2020). Viral sequences are often unstable in *Escherichia coli* and *Agrobacterium tumefaciens* due to prokaryotic promoter-like elements in the viral genomes and spontaneous insertions or deletions be a problem (Johansen, 1996; Lopez-Moya & Garcia, 2000). Rescue of plant rhabdoviruses has been achieved for Sonchus yellow net virus (SYNV) in *Nicotiana benthamiana* (Ganesan et al., 2013; Wang et al., 2015) and barley yellow striate mosaic virus (BYSMV) (Gao et al., 2019) and northern cereal mosaic virus (NCMV) in barley (Fang et al., 2022) by co-expressing RNPs and VSRs. More recently, reverse genetics were developed for segmented NSRs including tomato spotted wilt tospovirus (TSWV) (Feng et al., 2020), rose rosette virus (RRV) (Verchot et al., 2020), and a mini-replicon system for rice stripe tenuivirus (RSV) (Feng et al., 2021).

NSRs represent a significant technical advance for expression of heterologous proteins and RNAs in plants and an efficient NSR vector for maize has not been developed. Here, we used maize mosaic virus (MMV) as a model to develop the first infectious clone of an alphanucleorhabdovirus. The ability of MMV to infect maize and sorghum, two important crop plants, and the planthopper vector, made MMV an attractive candidate for developing an infectious clone and an expression vector. *Alphanucleorhabdovirus maydis*, is a typical member of the genus *Alphanucleorhabdovirus* and one of the best characterized plant rhabdoviruses (Ammar et al., 2009; Whitfield et al., 2018). MMV is transmitted in a persistent, propagative manner by its vector, *Peregrinus maidis*. The virus accumulates throughout insect development and has a broad tissue tropism in its vector (Ammar et al., 2009; Barandoc-Alviar et al., 2016; Yao et al., 2013). The MMV genome consists of 12,170 bases and encodes six proteins in the order 3’-N-P-3-M-G-L-5’ (Martin & Whitfield, 2019). Cellular localization and interactions of MMV proteins are well conserved between plant and insect cells (Martin and Whitfield, 2018). We stabilized the viral polymerase in *E. coli* and *Agrobacterium* using a plant intron. MMV is a versatile tool for stable expression of heterologous proteins in maize and planthoppers.

## 2. RESULTS

### 2.1 Insertion of plant intron in the L gene increased the stability of the MMV infectious clone

During construction of plasmids (pJL-L and pJL-MMV-WT) containing L gene sequences, we observed that initial transformed colonies contained the entire desired fragment upon colony-PCR with MMV N, P and L gene specific primers (Table S1). However, plasmid extracted from overnight bacterial culture produced a low quantity of pJL-MMV-WT and pJL-L plasmid. Moreover, none of the extracted plasmids generated desired fragments when validated with plasmid PCR and gene specific primers (Table S1). Growing bacterial cultures with plasmids at lower temperatures (25°C and 30°C) also did not solve the viral genome instability in *E. coli*. Instability of full-length MMV infectious clones and L gene in *E. coli* and *Agrobacterium* could be due to its large size or expression of toxic viral products from the MMV genome. We added an intron between the 3281 and 3282 splice sites (CTGCGGACAG^GTATCGATAT) of the L gene to improve the stability of pJL-L and pJL-MMV-WT constructs and to prevent the expression of polymerase gene that appear harmful to bacteria. Plasmid extracted from the overnight grown bacteria retained the full-length L with intron (Table S1). These results indicate that the presence of plant intron in the MMV L gene increased the stability of the plasmid and growth of *E. coli* and *Agrobacterium* in liquid cultures at different temperatures for both L by itself and when present in the L of the full-length viral genome.

### 2.2 Recovery of infectious MMV from cloned cDNAs in *N. benthamiana* leaves

To establish a reverse-genetics system, NSRs require in vivo reconstitution of active replicase complexes including the N, P and L proteins (Figure 1a). Thus, a mixture of *Agrobacterium* cultures harbouring the pJL-MMV-WT and the supporting plasmids, pTF-N&P and pJL-L-intron, were co-infiltrated into *N. benthamiana* leaves. No visible symptoms were observed in both infiltrated and systemic leaves at 12 days post infiltration (dpi) in replicated experiments. Anti-MMV virion immunoblots revealed viral proteins (MMV N, P, M, and G) in infiltrated but not systemic leaves (data not shown). This indicates that wild-type MMV failed to move systemically in infiltrated *N. benthamiana* leaves. To monitor virus infection *in vivo*, we engineered the MMV anti-genome with a GFP gene insertion between the duplicated N/P gene junction (Figure 1b). *N. benthamiana leaves* were agroinfiltrated with pJL-MMV-GFP together with expression constructs of N, P and L genes. At 14 dpi, GFP expression was observed in individual cells of infiltrated leaves of *N. benthamiana* (Figure 2a). GFP fluorescence was not observed in the pJL-MMV-GFP without expression constructs of N, P, and L (Figure 2a). Immunoblot analysis showed the evidence of viral proteins (MMV N, P, M and G) and GFP (Figure 2b) expression in the infiltrated leaves and indicates that they functionally interact with the agRNA transcripts to rescue recombinant MMV tagged with GFP (MMV-GFP) in the agroinfiltrated *N. benthamiana* leaves.

**Figure 1.**
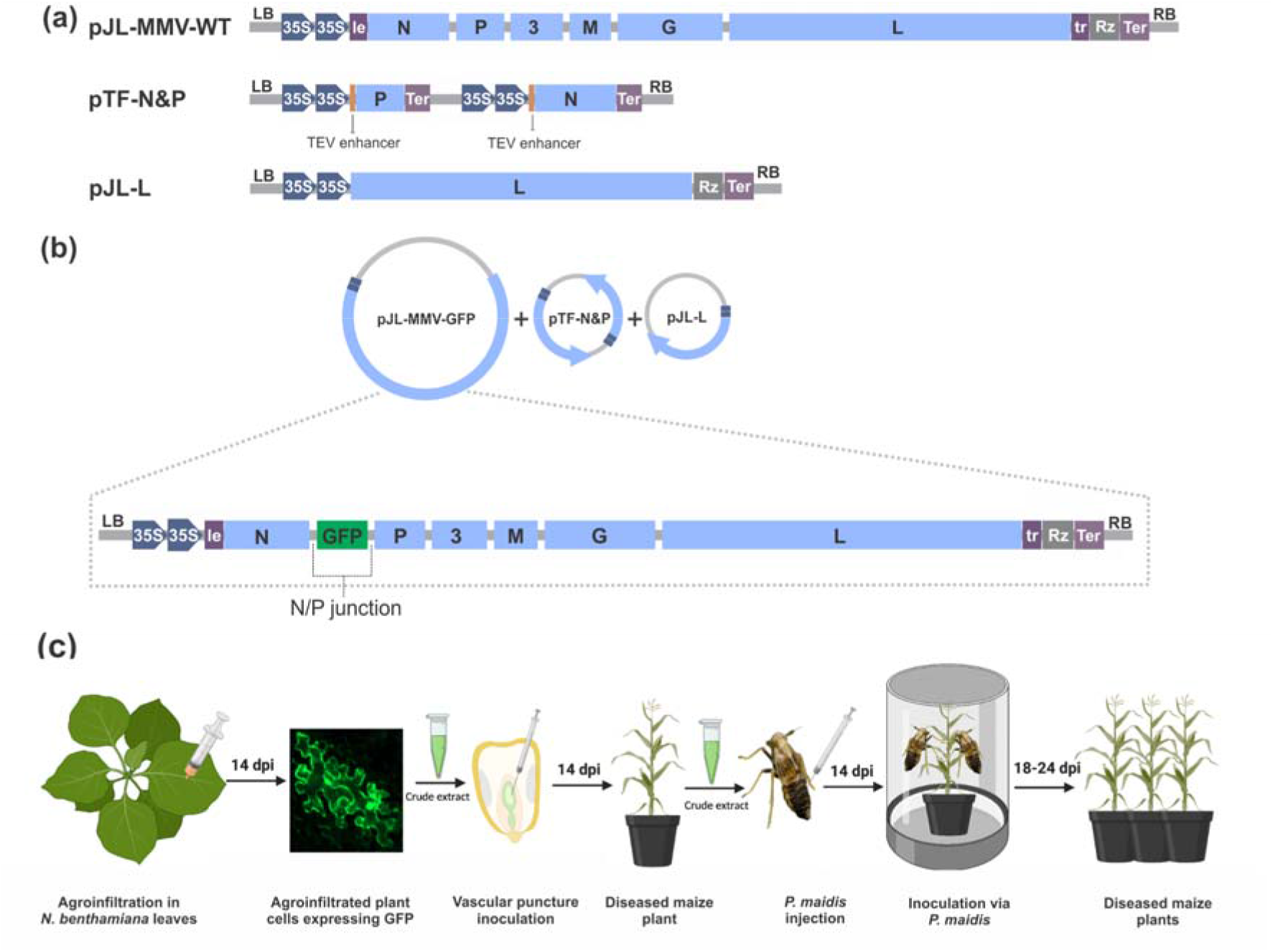
Rescue of infectious MMV from full-length cDNA clones in plants. a) Schematic diagrams of the pJL-MMV-WT, pTF-N&P and pJL-L-intron plasmids. The full-length MMV-plasmid designed for transcription to yield the MMV antigenome RNA (agRNA) and contains the full-length MMV cDNA positioned between a truncated CaMV double 35S promoter (2X35S) and the hepatitis delta ribozyme (Rz) sequence in pJL89 binary plasmid. Note that the sequences are shown in antigenomic (mRNA) sense. The full-length cDNA of N and P were inserted between the 2X35S and 35S terminator sequences in pTF binary plasmid. The full length cDNA of L with plant intron *ST-LS1* inserted in between 2X35S and 35S terminator sequences in pJL89 binary plasmid. b) Schematic representation of agroinfiltration with *Agrobacterium* strains containing pJL-MMV-GFP, pTF-N&P and pJL-L-intron plasmids and illustration of the pJL-MMV-GFP plasmid construction. The full length pJL-MMV-GFP contains duplicate N/P gene junctions flanking the GFP gene between the N and P genes of MMV antigenome cDNA. le: leader; tr: trailer; Ter: terminator; TEV: Tobacco etch virus; LB: left border sequence; RB: right border sequence. c) Illustration of the MMV rescue procedure and transfer to maize and planthoppers. dpi: days post inoculation. Figure 1c: created with BioRender.com.

**Figure 2.**
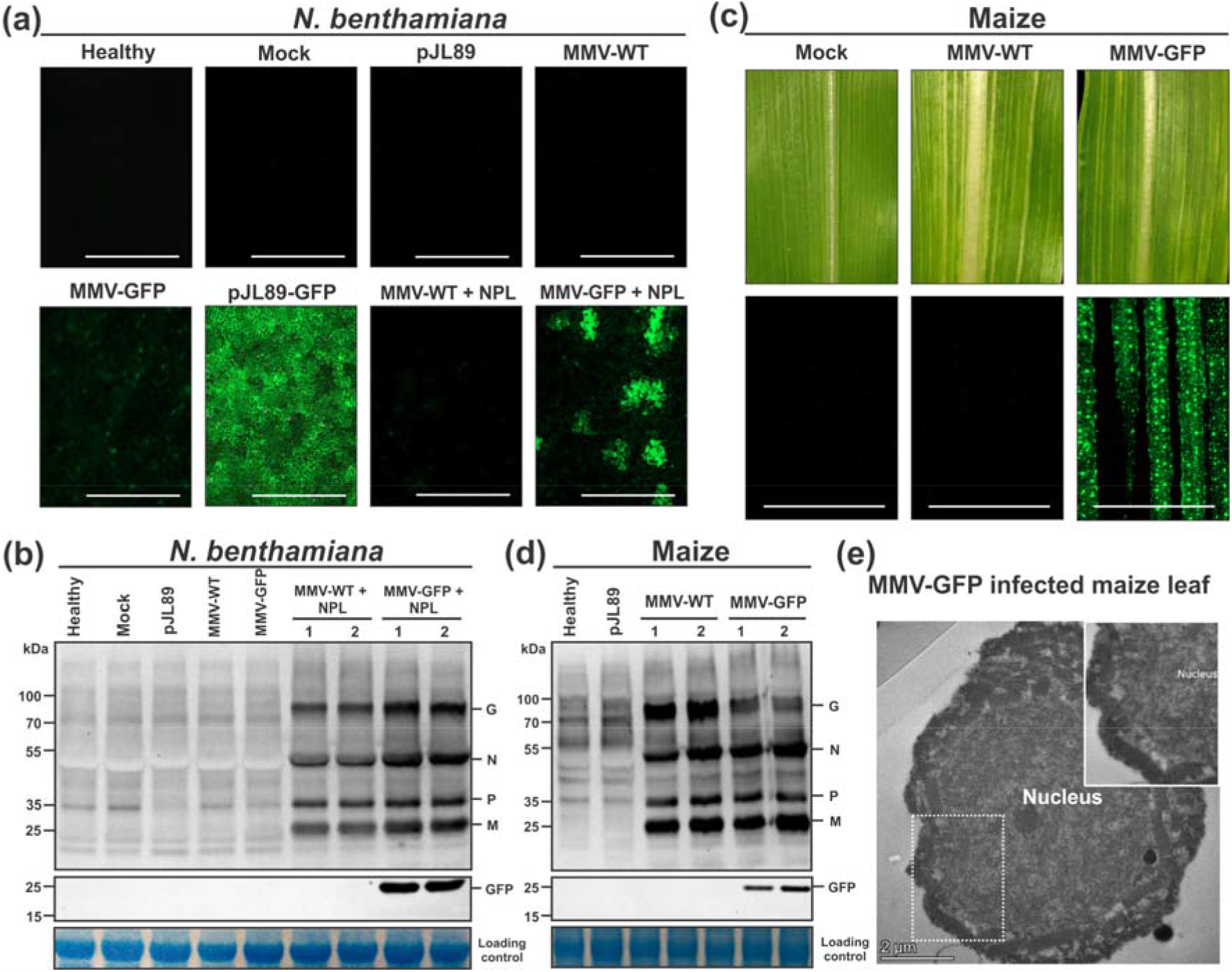
Maize mosaic virus (MMV) expression of GFP in the *N. benthamiana* and maize plants. a) Foci of GFP in *N. benthamiana* leaves after agroinfiltration with *Agrobacterium* strains containing mock (*Agrobacterium* alone), MMV-WT alone, MMV-GFP alone, pJL-GFP, MMV-WT+NPL and MMV-GFP+NPL. GFP foci were photographed at 14 days post infiltration (dpi) with a fluorescence microscope (Bars, 1000um). b) Immunoblot analysis of MMV and GFP proteins in *N. benthamiana* plants with the anti-MMV virion and anti-GFP antibodies, respectively. c) Maize leaves showing systemic mosaic symptoms 14 dpi with MMV-WT and MMV-GFP infected *N. benthamiana* sap by vascular puncture method. Images of maize leaves expressing GFP in the MMV-GFP infected plants. GFP foci were photographed at 24 dpi with a fluorescence microscope (Bars, 1000 µm). Mock plants inoculated with pJL-MMV-GFP alone crude extract did not show any symptoms. d) Immunoblot analysis of MMV and GFP proteins in maize plants with the anti-MMV virion and anti-GFP antibodies, respectively. GelCode Blue safe (ThermoFisher Scientific) stained Rubisco bands served as loading control. M: Prestained Protein Ladder (10-250kDa). e) Transmission electron micrograph of thin sanctions of MMV-GFP-infected maize showing particle accumulation in the nucleus. The dotted square box indicates the portion of the image that is magnified in the inset. Bars, 2 µm.

### 2.3 Rescue of MMV-WT and MMV-GFP in maize

To test the ability of the MMV infectious clone to infect maize plants, crude extracts obtained from *N. benthamiana* leaves agroinfiltrated with pJL-MMV-WT or pJL-MMV-GFP + NPL were inoculated to maize kernels by vascular puncture inoculation (VPI). After 14 dpi, pJL-MMV-GFP plants developed mosaic symptoms on systemic leaves, similarly to those observed in wildtype-MMV infections. In addition, MMV-GFP + NPL resulted in high-intensity GFP fluorescence that was systematically distributed in the symptomatic young leaves (Figure 2c). Immunoblot analysis confirmed that MMV viral proteins (N, P, M and G) and GFP accumulated in systemically infected maize leaves (Figure 2d). Abundant bacilliform particles were observed in the nucleus of thin sectioned cells infected with pJL-MMV-GFP (Figure 2e).

### 2.4 Stable GFP expression by MMV vector in maize and planthoppers following virus passages

We demonstrated that the MMV-GFP vector can be successfully used to stably express the GFP in passages between various hosts: from *N. benthamiana* to maize; maize to maize; maize to *P. maidis;* and *P. maidis* to maize. Stability of the inserted GFP gene was evaluated in plants and planthoppers following three successive passages by RT-PCR using MMV N and GFP specific primers (Table S1). Furthermore, GFP fluorescence in the MMV-infected plants were monitored weekly (Figure 3 a,b). MMV-GFP infected leaves (5L-9L) from the base of the plant were tested for virus and GFP. The result showed presence of MMV N and full-length GFP bands in all the infected plants (Figure 3c,d). We also observed GFP in the MMV-GFP infected plant leaves using fluorescence microscopy for the lifetime of the plant (data not shown). When *P. maidis* adults were injected with MMV-GFP infected maize crude extract, 3 of 10 insects in the first passage, 4 of 10 in the second passage, and 4 of 10 in the third passage tested positive for MMV and GFP (Figure 3e). Only MMV-GFP infected insects tested positive for both MMV and GFP. After the 24 day inoculation access period (IAP), the infected *P. maidis* adults and nymphs showed GFP fluorescence (Figure 4a). Immunoblot analysis confirmed that MMV viral proteins (N, P, M and G) and GFP accumulated in systemically infected *P. maidis* adults. The presence of MMV N and full-length MMV GFP bands present in all the three passage generations in both plants and planthoppers showed insert stability in the MMV vector (Figure 3c-f). We observed that purified MMV-GFP virions injected to *P. maidis* nymphs showed GFP fluorescence from 7 dpi (Figure 4c). Moreover, we also used frozen MMV-GFP infected leaf crude extract for injections which also resulted in successful virus infection in insects and virus transmission.

**Figure 3.**
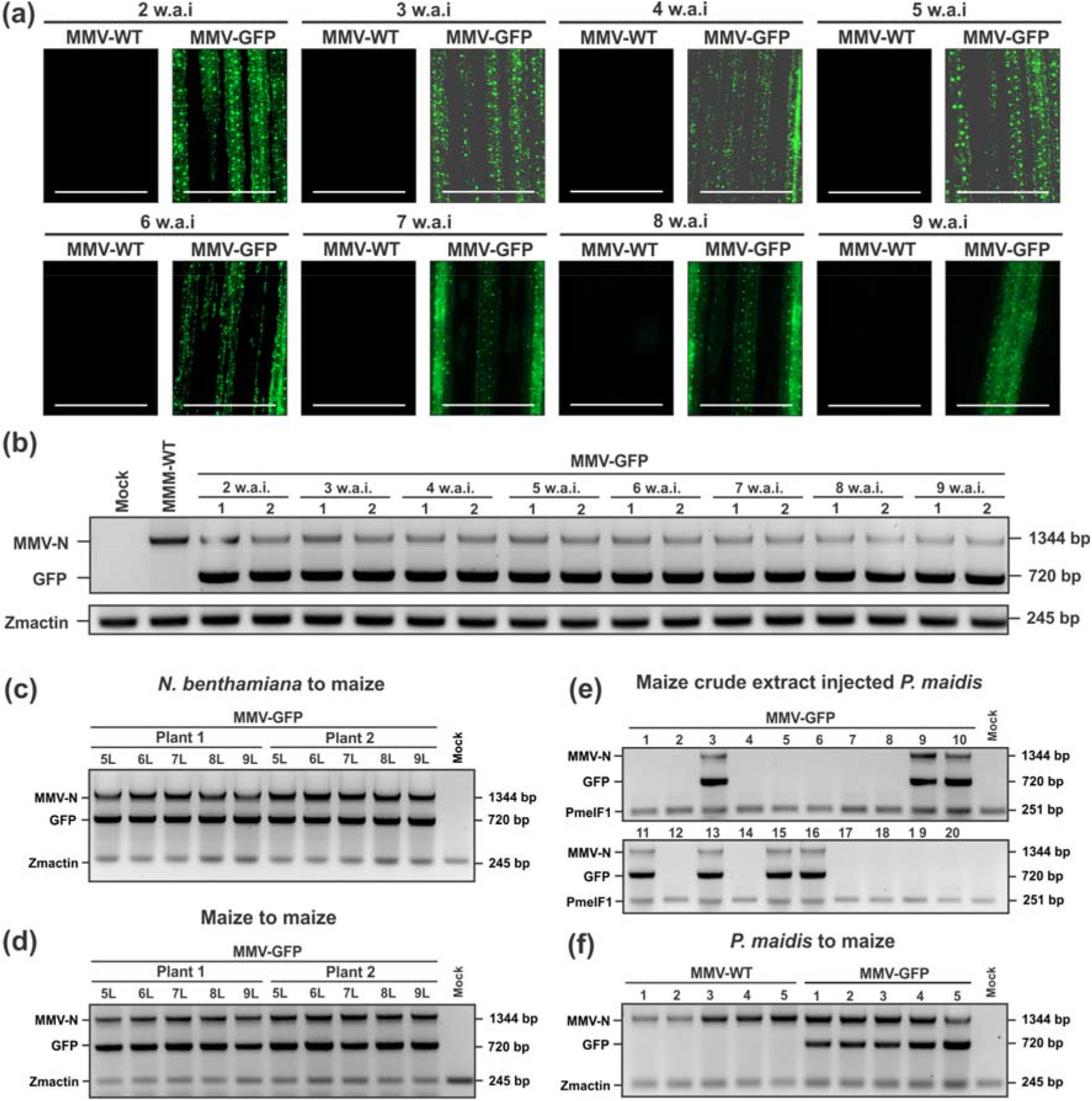
Time course observation of GFP expression in MMV-GFP infected maize leaves. a) GFP foci were photographed weekly at two to nine weeks after inoculation (w.a.i) with a fluorescence microscope (Bars, 1000 µm). Maize plants inoculated with MMV-WT did not show green fluorescence. b) RT-PCR analysis of virus and GFP in the MMV-GFP infected maize plants collected from 2nd to 9th w.a.i. c-f) Stability of GFP sequence carried by MMV vectors in all the three passage generations in both plants and planthoppers. c) RT-PCR analysis of maize plants inoculated with *N. benthamiana* crude extract harbouring MMV-GFP derivatives via VPI. d) RT-PCR analysis of maize plants inoculated with MMV-GFP infected crude extract via VPI. 5L, 6L, 7L, 8L and 9L indicate the leaf number that was sampled from the base of the plant. Plant 1 and Plant 2 indicate two individual MMV-GFP infected plants. e) RT-PCR analysis of individual *P. maidis* injected with MMV-GFP infected Rescue of an Alphanucleorhabdovirus infectious clone maize crude extract. f) RT-PCR analysis of MMV-GFP infected independent maize plants transmitted by *P. maidis*. Controls: Mock (Maize: *Agrobacterium* alone inoculated plants; *P. miadis*: healthy maize crude extract injected planthoppers) and MMV-WT: maize plants infected with MMV-wild type. MMV N and GFP primers to detect the virus presence and stability of the GFP insertion. *Zmactin* and *PmeIF*-1 were used as internal controls for maize and planthoppers respectively.

**Figure 4.**
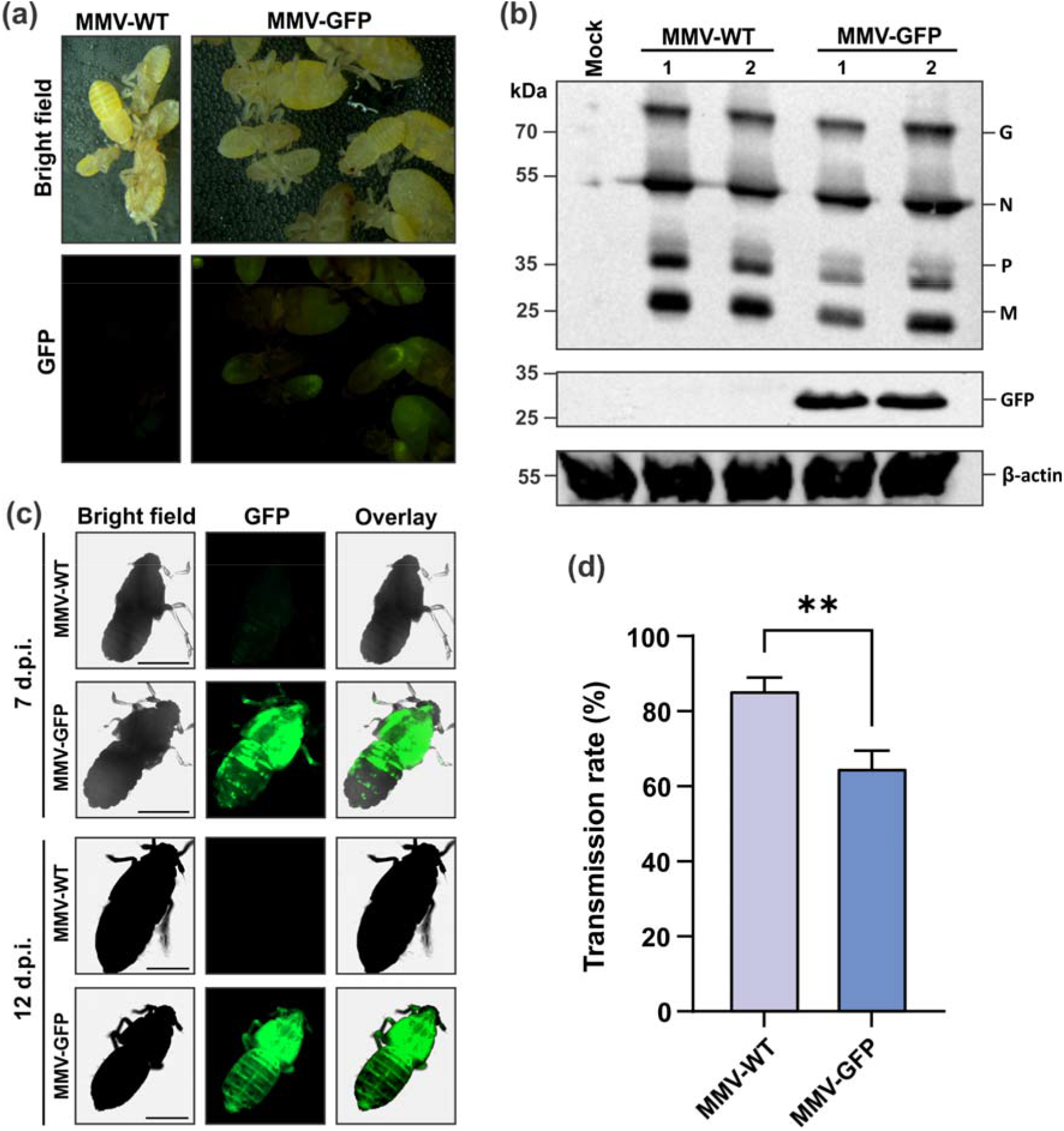
MMV expression of GFP in planthoppers and virus transmission efficiency. a) *P. maidis* adults and nymphs that fed on MMV-WT and MMV-GFP infected maize plants at 28 dpi were photographed with a stereomicroscope equipped with a GFP filterset. b) Immunoblot analysis of the expression of MMV and GFP expression in *P. maidis* adults infected with MMV and MMV-GFP. Total proteins were extracted from planthoppers that fed on healthy (mock) or MMV-WT and MMV-GFP infected maize plants were analyzed by immunoblotting with antibodies against whole MMV virion and GFP antibodies. Each lane represents a pool of 10 insects feeding on MMV-WT and MMV-GFP infected plants and the numbers 1 and 2 indicate independent samples for the respective treatments. GelCode Blue safe (ThermoFisher Scientific) stained protein bands served as loading controls. M: Prestained Protein Ladder (10-250kDa). c) MMV-GFP virions injected *P. maidis* nymphs and adults at 7 and 12 dpi were photographed with a fluorescence microscope (Bars, 1000 µm). d) Transmission of MMV-WT and MMV-GFP to maize plants by *P. maidis*. Forty nanoliters of MMV-WT and MMV-GFP infected plant crude extracts were injected into *P. maidis* nymphs. Seven days after the incubation period, insects were transferred to one-week old healthy maize plants for a two week inoculation access period. Each bar represents the mean and standard error (SE) of three experimental replicates, and each replicate consists of a group of 20 plants. Different letters represent differences in transmission between treatments (t-test, p < 0.01).

### 2.5 MMV-WT and MMV-GFP virus transmission

Transmission efficiency of MMV-WT and MMV-GFP was measured in experiments with *P. maidis* nymphs that were injected with MMV-WT or MMV-GFP crude extract from infected maize plants. For MMV-WT, after 14 days post inoculation access period, 19 out of 20 (95%) plants became infected in the first experiment, whereas, 18 out of 20 (90%) and 19 out of 20 (95%) plants developed mosaic symptoms on systemic leaves in the second and third experiments, respectively (Figure 4d). For MMV-GFP, after a 14 days post inoculation access period, 12 out of 20 (60%) plants in the first experiment were infected with MMV while 13 out of 20 (65%) and 14 out of 20 (70%) plants in the second and third experiments developed mosaic symptoms on systemic leaves, respectively (Figure 4d). Moreover, MMV-GFP plants showed delayed systemic symptom development compared to MMV-WT plants, these results suggested that MMV-GFP is replicating slowly after transmission. Insect vector survival and transmission efficiency were improved by injecting insects with infected maize extracts (Table S2). These results demonstrate that MMV-GFP rescued in inoculated maize plants by VPI could be transmitted to *P. maidis* by injection of leaf extracts and recombinant MMV-GFP is insect transmissible.

### 2.6 MMV gene expression does not differ between a MMV-WT and a MMV-GFP *in planta*

We evaluated the transcription levels using qRT-PCR of all MMV genes between MMV-WT and MMV-GFP infectious clones to verify the effect of introducing cargo proteins into the MMV infectious clone on the expression of downstream viral genes. For each gene, two sets of primers were used with similar efficiency (Table S1). The results demonstrated no significant effect on the gene expression of downstream genes in MMV genome when GFP was inserted at 3’ region of the genome (Figure 1b), as well as, no significant difference observed in viral gene expression between MMV-GFP and MMV-WT for both primers tested for each gene (*p*>0.05; Figure 5). Our results suggest that adding one single gene at the 3’ end of MMV genome did not affect downstream viral gene expression. However, the effect of multiple insertions and location of insertions in the MMV genome needs further characterization.

**Figure 5.**
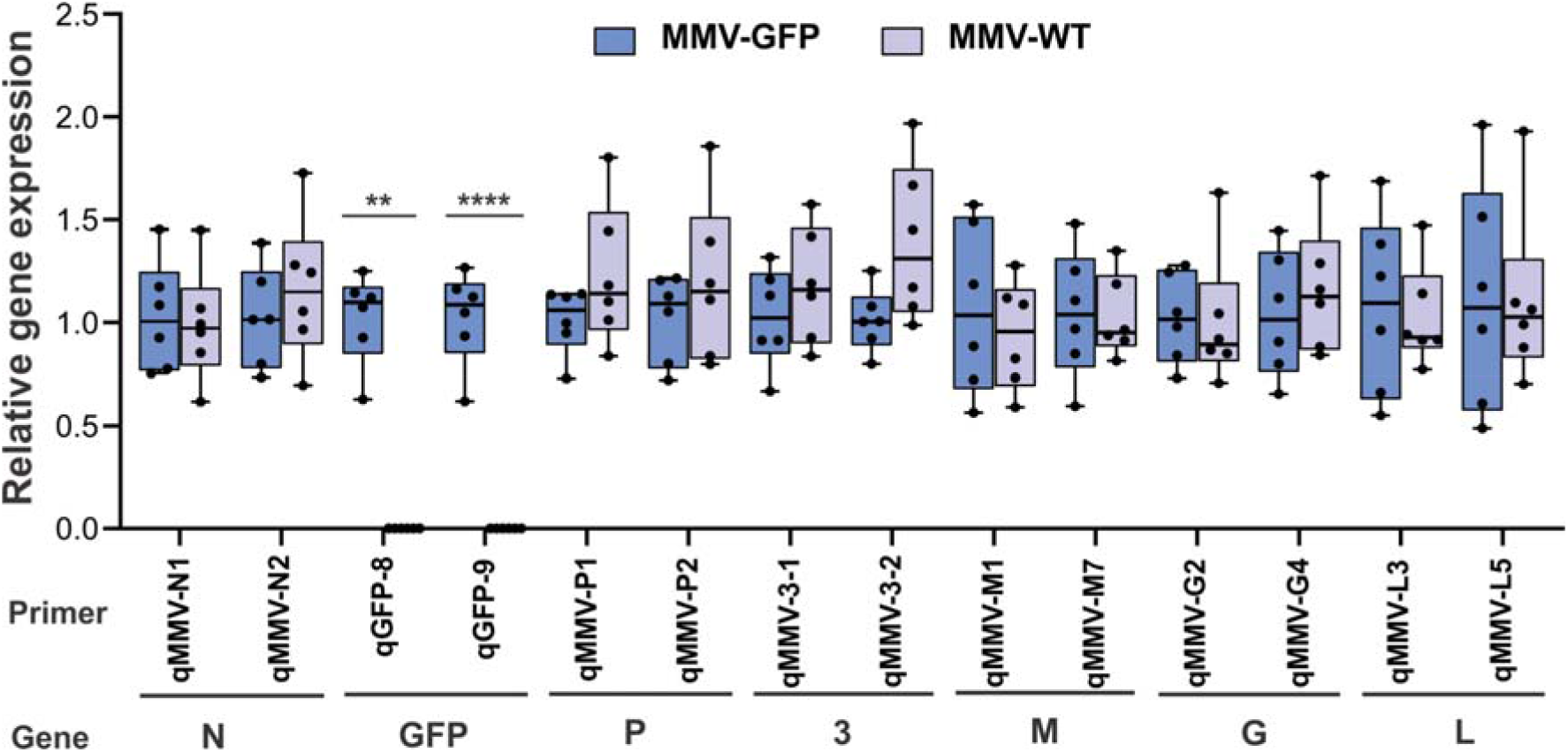
MMV gene expression did not differ between MMV-WT and MMV-GFP. Boxplots of relative viral gene expression comparing MMV-WT and MMV-GFP using two different pairs of primers for each gene (Table S1) are shown. Analysis was performed 30 days after plant inoculation. Boxplots represent minimum and maximum, median, and interquartile ranges of virus gene expression. Dots represent individual values from each biological replicate (n=6). No significant difference in gene expression was observed between MMV-WT and MMV-GFP (*p*>0.05) except for GFP (\*\**p*<0.01; \*\*\*\**p*<0.001), according to non-parametric Mann Whitney test. N; nucleocapsid protein; GFP: green fluorescent protein; P: phosphoprotein; 3: movement protein; M: matrix protein; G: glycoprotein and L: polymerase.

## 3. DISCUSSION

A reverse genetics system for plant NSRs viruses provides a platform for studying the basic biology of plant rhabdoviruses and for delivery of RNAs and proteins to crop plants with the goal of characterizing and improving agronomic traits. In addition, plant virus-based expression vectors have been used for the expression of vaccine antigens, antibodies and other therapeutic proteins (LeBlanc et al., 2021). To date, a few maize viruses have been developed for transient gene function studies (reviewed in Kant and Dasgupta, 2019; Xie et al., 2021, Moltshwa et al., 2020). While important, these systems have limited use for maize researchers, due to transgene instability, narrow host range in monocots, and severe symptoms in inoculated plants (Ding et al., 2006; van der Linde et al., 2011; Benavente et al., 2012). Improvements in high-yielding hybrid crops, including maize, through transgenic modification is complex, and is limited to a few genotypes, requiring multiple generations to produce agronomically viable lines. In contrast, virus delivery technologies could hasten the breeding process by enabling direct modification of advanced breeding line varieties with desirable traits. Here, we developed an efficient NSR viral vector for expressing heterologous proteins in maize.

The creation of infectious clones for plant NSRs continues to be technically challenging with each system requiring a unique suite of “tricks” to make it work (German et al., 2020). During the construction of a MMV infectious clone, we observed plasmid instability because of spontaneous deletions of viral polymerase sequences during propagation of the cDNA clones in *E. coli* and *A. tumefaciens* cells. To circumvent the problem of instability of plasmids that contained the L gene, we introduced a plant intron into the polymerase gene L. This insertion increased the stability of plasmids during propagation in *E. coli* and *Agrobacterium*. Previous studies showed that introduction of a plant intron into the plant virus infectious clones has proven to be an efficient way to increase stability for viruses belonging to the genera *Tobravirus* (Ratcliff et al., 2001) and *Potyvirus* (Johansen, 1996; López-Moya & Garcia, 2000; Sun et al., 2017; Tran et al., 2019). Alternatively, Feng et al. (2020) used a codon optimization strategy to remove potential splice sites in the TSWV polymerase sequence. However, codon optimization can lead to alterations in protein conformation and function (Mauro & Chappell, 2018). Codon optimization was not required for successful reconstitution of MMV and that may be partly due to the biology of nuclear-replicating viruses like MMV because their mRNAs are adapted for nuclear exit similar to host mRNAs. We anticipate the introduction of introns for increasing L gene stability can be adapted to construct other plant rhabdovirus infectious clones. In this study, MMV was rescued in *N. benthamiana* without VSRs, although VSRs were required for the initial rescue of other plant rhabdoviruses (SYNV, BYSMV, and NCMV) to increase the accumulation of core proteins and minireplicon derivatives (Wang et al., 2015, Gao et al., 2019, Ma & Li, 2020, Fang et al., 2022).

Rescue of infectious plant NSR viruses from cDNA strictly requires agrobacterium mediated delivery of core replication proteins together with their genome RNAs. However, it is a major challenge to deliver multiple essential components into maize cells via agroinoculation/biolistic delivery. Our attempts to rescue MMV infectious clones via agroinoculation did not achieve systemic infection in multiple attempts. The main obstacle is the lack of efficient gene expression systems to rescue monocot-infecting rhabdoviruses. Hence, for the first time we used the VPI method to move MMV from *N. benthamiana* to maize. Though we observed a low rate of infection via VPI (Table S2), once we moved MMV from *N. benthamiana* to maize, we could efficiently generate virus infected plants via insect transmission.

Similar to the work with BYSMV (Gao et., 2019) and NCMV (Fang et al., 2022), monocot-infecting cytorhabdoviruses, MMV replicated in initial *N. benthamiana* cells that received the ectopically expressed viral replicase proteins (N, P, and L) and gRNA but was unable to move cell-to-cell or systemically. In contrast, SYNV, the first plant rhabdovirus infectious clone, is able to systemically infect *N. benthamiana* and this feature of SYNV biology facilitated development of this virus as a vector (Wang et al., 2015; Peng et al., 2021). The addition of GFP to the MMV cDNA clone between N and P genes with duplicated N/P gene junctions enabled tracking of virus infection in plants and insects. We excised GFP expressing cells harbouring recombinant MMV virions in the agroinfiltrated leaves and injected cell extract to maize seeds using the VPI method. This allowed us to rescue the first alphanucleorhabdovirus entirely from cDNA clones and to introduce the virus to its natural host, maize. Inoculation of maize via agroinoculation or biolistic delivery of plasmids was unsuccessful for both MMV-WT and MMV-GFP (data not shown). Initial attempts to infect *P. maidis* by injecting *N. benthamiana* crude extract harbouring MMV-WT did not succeed in transferring infectious virus from *N. benthamiana* to the insect vector. It is likely that the secondary metabolites in the crude extracts of tobacco were toxic to the injected planthoppers.

The MMV infectious clone is a tractable system for stable expression of foreign genes in maize and insects over repeated passages. We observed that recombinant MMV could be stored as infected tissue or purified virus at -80°C and revived in maize by VPI or insects by microinjection. This shows that recombinant MMV can be easily maintained long term. Comparison of gene expression between MMV-WT and MMV-GFP revealed that there were no significant differences in relative abundance of MMV transcripts downstream of the GFP insertion in the MMV genome. This suggests that even with the additional ORF and gene junction sequences the expression of MMV genes and/or stability of transcripts remains unchanged. The development of the BYSMV infectious clone was a major step forward for NSR viral vectors of monocots including barley, wheat, and foxtail millet; however, BYSMV distribution in maize leaves appeared patchy (Gao et al., 2019). The MMV clone supported strong expression of GFP throughout the entire maize leaves over eight weeks post inoculation. The development of an efficient NSR vector for maize overcomes some of the drawbacks associated with positive strand RNA virus (+ssRNA)-based tools that are currently available for monocots such as insertion size limits and insert instability during passages (Mei et al., 2019; Bouton et al., 2018; Cheuk and Houde, 2018, Jarugula et al., 2018; Tatineni et al., 2011). The differences in insert stability are due to the high recombination rates reported for +ssRNA viruses and the low to no recombination rates observed for NSR genomes (Chare et al., 2003, Patiño-Galindo and Rabadan, 2021). The stability of the insertions in the MMV genome make it a good candidate viral vector for expression of genes for crop improvement because they can be expressed over a long period of plant growth in the field or greenhouse. Similar stability and expression characteristics for MMV were observed in maize and planthopper vectors.

Here, we have developed a new virus vector using the maize alphanucleorhabdovirus, MMV, and demonstrated the successful GFP expression in both plants and insects vector *P. maidis*. The major advantage of rhabdoviruses is that they are able to replicate in both plants and insect vectors, providing a convenient and highly efficient approach for transmission of gene editing reagents to multiple hosts through virus infection and insect transmission. Previous studies of MMV transmission biology, molecular interactions between *P. maidis* and the virus, and genome editing in the vector make this system a good platform for further characterizing rhabdovirus-vector interactions (Barandoc-Alviar et al., 2016; Barandoc-Alviar et al., 2022; Klobasa et al., 2021). Other rhabdoviruses, BYSMV (Gao et al., 2019) and SYNV (Ma et al., 2020), can express multiple guide RNAs and Cas nucleases. Thus far, successful genome editing has been demonstrated in *N. benthamiana* using rhabdovirus vectors but with the ease of manipulation of MMV after initial launch, we expect to see rhabdovirus-based editing in important crop plants in the near future as this is now a possibility for maize. Creation of an MMV infectious clone opens the door to a wide range of applications for expression of foreign genes and guide RNAs in maize and insects and for genome editing to enhance crop productivity.

## 4. EXPERIMENTAL PROCEDURES

### 4.1 Construction of MMV infectious clone and introduction of intron into L gene

The full-length MMV (12,170 bp; accession number MK828539) cDNA was synthesized in pcc1-pbrick plasmid between cauliflower mosaic virus promoter (CaMV) 35S promoter and CaMV PolyA terminator *de novo* by GenScript, Piscataway, New Jersey. To assemble full-length MMV-WT from the entire cDNA into pJL89 binary plasmid (Lindbo, 2007), fragments of 12,205 bp and 4137 bp were amplified from pbrick-MMV and pJL89 using pJL-MMV F/R and pJL F/R primers, respectively (Table S1). The forward primer contains 17 nt overhangs complementary to the 3’ end of the double 35S promoter, whereas reverse primer contains 18 nt overhangs complementary to the 5’ end of the self-cleaving hepatitis delta virus ribozyme (Rz), respectively. The two overlapping fragments were amplified using Q5 high-fidelity DNA polymerase (NEB) and introduced into the pJL89 vector by one-step NEBuilder Hifi assembly (NEB) (Figure 1a). To generate pTF-N&P (plasmid for expression of N and P proteins) (Frame et al., 2002), MMV N and P were amplified using MMV NF/NR and PF/PR primers from cDNAs synthesized from infected-plant RNA and subcloned into pTF binary plasmid between the double CaMV 35S promoter and nopaline synthase (NOS) terminator using Gateway Clonase enzyme mix (Invitrogen) (Figure 1a). To generate pJL-L for expression of the polymerase, two overlapping PCR fragments 6936 bp and 4137 bp were amplified from pbrick-MMV and pJL89 using 35SLF/TerLR and pJL F&R primers, respectively (Figure 1a). Competent cells of *E. coli* strain Top 10, DH10B, DH5α (ThermoFisher Scientific) and NEB Stable Competent *E. coli* (NEB) were used for cloning steps. Except pTF-N&P, all the competent cells mentioned above tended to lose the full-length MMV and L gene plasmids in overnight cultures at three different temperatures (25°C, 30°C and 37°C).

To increase the stability of the plasmid, intron 2 (189 bp) of the light-inducible gene ST-LS1 (X04753) from *Solanum tuberosum* was introduced into the L gene between 3281 and 3282 splice site (CTGCGGACAG^GTATCGATAT) nucleotides (Table S3). The putative intron splicing sites of wild type L gene sequences was predicted by Alternative Splice Site Predictor (ASSP) (Wang & Marin, 2006). To generate pUC-MMV-L-intron (6784 bp), the MMV L gene from 2893 nucleotides-*ST-LS1*-pJL89 terminator in pUC57 was synthesized de *novo* by GenScript. To generate pJL-L-intron, three overlapping PCR fragments LF1 (2,943 bp) and LF2 (3,487 bp) and a third fragment pJL89 (LF3, 4,137 bp) were amplified from pbrick-MMV, pUC-MMV-L-intron and pJL89 plasmids (Figure 1a) with the primer pairs 35SLF/MMVLXSR; LXSF/TerLR and TerLF/pJLR respectively. To generate pJL-MMV-intron, three overlapping PCR fragments of MF1 (3,489 bp), MF2 (9,224 bp) and MF3 (4,137 bp) were amplified from pUC-MMV-L-intron, pbrick-MMV and pJL89 plasmids, using primer pairs LXSF/TerLF; pJLMMVF/MMVLXSR and TerLF/pJLR, respectively. The overlapping fragments were purified and further assembled using NEBuilder Hifi assembly (NEB). The resulting pJL-L-intron was cloned in a binary vector pJL89 between the double 35S promoter (2X35S) at 5’ terminus and the HDV ribozyme at the 3’ terminus. All the fragments (MF1, MF2 and MF3) were amplified using Q5 high-fidelity polymerase (NEB) and introduced into binary plasmids using NEBuilder Hifi assembly (NEB) (Figure 1a). When the *E. coli* transformants were cultured overnight, pJL-L and pJL-MMV-WT with the addition of the *ST-LS1* intron resulted in a single plasmid with expected characteristics when analyzed by PCR and restriction digestion. Primers used for NEBuilder Hifi assembly (NEB) contained 10-15nt overhangs at the 5’ end that was homologous to the 5’ end of another PCR product, so that the two PCR products could be ligated and circularized. All PCRs were carried out using Q5 high fidelity polymerase (NEB). NEB Stable Competent *E. coli* (NEB) were used in these cloning steps. All the primers used in this study were listed in Table S1.

To develop a recombinant MMV vector for foreign gene expression in plants, a duplicated N/P gene junction sequence along with the enhanced green fluorescent protein (eGFP) coding sequence were inserted into the pJL-MMV-WT plasmid between the N and P genes to generate pJL-MMV-GFP. The complete coding region of GFP followed by the N/P gene junction sequence, was synthesised by GenScript. To facilitate sub-cloning, an Bsu36I (ThermoFisher Scientific) site was introduced at either ends of the GFP and N/P gene junction clone. Both pJL-MMV-WT and pUC57-GFP-N/P plasmids were digested by *Bsu*36I and ligated to generate pJL-MMV-GFP (Figure 1b). In this configuration, GFP mRNA synthesis is initiated immediately after termination of the upstream N protein mRNA synthesis by the duplicated N/P gene junction and is followed by P mRNA synthesis that is directed by the native N/P gene junction. Clones with correctly oriented GFP-N/P inserts were verified by MMV1355F and MMVPR primers and sequencing. All constructs were confirmed by sequencing and transformed into *A. tumefaciens* GV3101.

### 4.2 *Agrobacterium* infiltration

Virus infection of *N. benthamiana* was achieved by *Agrobacterium*-mediated transient gene expression of infectious constructs from the T-DNA of a binary plasmids pJL89 and pTF. *A. tumefaciens* GV3101 with pJL-MMV-WT, pJL-MMV-GFP, pJL-L plasmids (Kanamycin, 50 µg/ml) and pTF-N&P plasmid (Spectinomycin, 50 µg/ml) were individually grown in Luria-broth containing rifampicin (50 µg/ml) and Gentamicin (50 µg/ml). Cells grown overnight were harvested by centrifugation and resuspended in the infiltration buffer and incubated for 3h at room temperature in the presence of 100 µM acetosyringone, as described in Kanakala et al. (2019). After incubation, equal volumes of *Agrobacterium* cultures harbouring the pJL-MMV-WT or pJL-MMV-GFP alone and pJL-MMV-WT or pJL-MMV-GFP + NPL were co-infiltrated into the *N. benthamiana* with a final OD_600_ of 0.5 for each construct.

### 4.3 Vascular puncture inoculation, crude sap-injections and insect transmission studies

*N. benthamiana* leaf extracts infiltrated with pJL-MMV-WT and pJL-MMV-GFP plasmids were homogenised in extraction buffer (10mM Potassium buffer; pH 7) and centrifuged at 12, 000 g for 5 min at 4°C (Louie, 1995). Then 5 µl of crude supernatants were vascular puncture inoculated using a 5 pin-tattoo needle into the overnight soaked maize kernels at 30°C (Figure 1c). Then the VPI seeds were incubated in a humid tray at 30°C for two days before sowing. These plants were subsequently observed for systemic symptom expression or image analysis. The symptomatic plant leaves showing mosaic symptoms and GFP expression were homogenized in extraction buffer (100mM Tris-HCl, 10mM Mg (CH3COO)2, 1mM MnCl_2_, and 40mM Na_2_SO_3_, pH 8.4) and centrifuged at 12 000 g for 10 min at 4°C (Jackson & Wagner, 1998). Then, 40 nl of the crude extract supernatants were injected into the anesthetized *P. maidis* (nymphs) using a Nanoinject III Programmable Nanoliter injector (Drummond Scientific Co.) (Figure 1c). In three independent experiments, the surviving injected nymphs were maintained on the healthy maize seedlings for a 7 days incubation period and then transferred to healthy maize seedlings for a 2-week IAP (Figure 1c). Adult insects injected with extraction buffer alone (∼30/replicate) were used for control experiments. Symptoms in plants and GFP expression in systemic leaves and insects were observed and recorded weekly after infection.

### 4.4 Electron microscopy and Image acquisition

Systemically infected leaf tissues were fixed in 2.5% glutaraldehyde and 1% osmium tetroxide (both in 100mM sodium cacodylate buffer, pH 7.0) as previously described Kong et al., 2014. Following ethanol dehydration, the fixed tissues were then embedded in Spurr’s resin (Spurr, 1969) as instructed by the manufacturer (Sigma-Aldrich). Ultrathin sections (90nm) were cut with a Diatome diamond knife from the embedded tissues using the Leica EM U7 Ultramicrotome (Leica Microsystems). The thin sections were then stained with 4% uranyl acetate for 20 minutes followed by lead citrate for 30 seconds. The stained sections were imaged on a ThermoFisher Talos F200X TEM operated at 80 KV to determine virion structure in the infected cells.

The fluorescence of plant leaves and insects was observed with the objective of 4x using the BioTek slide reader and Image Software, version 3.04. Images were captured using default settings. Leaves agroinfiltrated with pJL89 alone were used as a negative control.

### 4.5 RNA extraction, cDNA synthesis and RT-PCR and quantitative real-time PCR (qRT-PCR)

To assess reporter gene stability, multiple successive passage experiments were performed in the following order: 1) *N. benthamiana* to maize, 2) maize to maize, 3) maize to *P. maidis* and 4) *P. maidis* to maize. During *N. benthamiana* to maize and maize to maize passages, we collected infected leaves from 2 to 9 weeks after inoculation (w.a.i.) for further analysis. To determine the passage of maize to *P. maidis*, MMV-GFP infected crude sap was injected into insects and collected two weeks after release on maize plants for further analysis. To determine virus passage from *P. maidis* to maize, we collected infected maize leaves for further analysis. Leaves of vascular puncture inoculated maize plants and crude sap injected adult *P. maidis* were harvested for RNA extraction using the RNeasy Plant Mini kit (Qiagen) and TRIzol reagent (ThermoFisher Scientific) according to the manufacturer’s recommendations, respectively. For virus detection and gene expression quantification, an aliquot containing 1.5 microgram of total RNA was used for the first strand cDNA synthesis using the Verso cDNA Synthesis Kit with RT-enhancer (ThermoFisher Scientific) to remove residual genomic DNA, following the manufacturer’s instructions. After first strand cDNA synthesis, primer pairs MMV N F&R, GFP F&R were used to detect the presence of MMV by RT-PCR. *Zea mays actin* and *P. maidis eIF1* genes were used as an internal control with primer pairs *Zmactin* F&R and *PmeIF1* F&R respectively.

To compare gene expression between MMV-WT and MMV-GFP *in planta*, single maize plants were inoculated by a single viruliferous *P. maidis* males brachypterous from an age calibrated colony according to Yao et. al., 2019. Thirty days after inoculation, the second youngest maize leaf was collected from symptomatic plants and 50 mg of tissue was subjected for RNA extractions and used for cDNA synthesis. Primers were designed for all MMV genes using the PrimerQuest Tool software from Integrated DNA Technology (IDT) using as template the full-length MMV genome (GenBank accession number MK828539), with default settings. A total of 34 pairs of primers were first tested for efficiency by using five-fold serial dilutions of cDNA synthesized using oligo-dT from total RNA obtained from MMV-GFP infected maize plants as template for qRT-PCR (Table S1). Efficiency was calculated based on the standard curve method. First, primer pairs with efficiencies between 90% and 110% and showing a single peak based on the melting curve were selected for further analyses. The highest difference in efficiency between selected primers pairs for a given gene was 6% (Table S1). In a final volume of 10 μl, the qRT-PCR reactions contained 4 μl of 7-fold diluted cDNA, 1 μl of mixed primers containing (5 M of each primer (forward and reverse), and 5 μl of iTaq Universal SYBR Green Supermix (BioRad). The qRT-PCR cycles consisted of an initial denaturing step at 95°C for 1 min, followed by 40 cycles of 95°C for 15 sec and 60°C for 1 min followed by melting curve analyses. All reactions were performed in duplicates. Expression levels for each gene were quantified by the comparative cycle threshold method (2^-ΔΔCT^) (Livak and Schmittgen, 2001) using two primers pairs for each gene and the membrane protein (MEP) as internal reference gene (Manoli et al., 2012) (Table S1). Statistical analyses were performed using GraphPad Prism software version 9.2.0. To compare differences in gene expression between MMV-WT and MMV-GFP the expression data were log_10_ transformed and subjected to the non-parametric Mann Whitney test.

### 4.6 Immunoblot analysis of viral proteins

For immunoblot analysis, proteins were separated on 4-15% gradient Mini-PROTEAN TGX Precast gels (Bio-Rad). Immunoblot analysis of the plant and insect protein extracts was performed according to the procedures described previously (Caplan et al., 2008). Proteins were transferred to nitrocellulose membranes using the Trans-Blot Turbo Transfer system (Bio-Rad). Membranes were blocked using 5% non-fat dry milk for 1 h then washed in 3x in Tris-buffered saline containing 0.1% Tween-20 (TBS-T). Transferred membranes were incubated with anti-MMV virion (made against purified MMV virus), this antibody recognizes the main structural proteins of the virus (N, P, M, and G when used at 1:2,000 dilution) and anti-GFP conjugated with horseradish peroxidase (Invitrogen, 1:2,000 dilution) with gentle shaking overnight at 4°C to detect MMV proteins and GFP respectively. Then the antibody incubated membranes were washed 3 times with TBS-T before adding secondary antibody goat anti-rabbit (whole virion) IgG-conjugated alkaline phosphatase for 2 h at room temperature with gentle shaking, then washed 3 times with TBS-T before addition of SuperSignal^™^ West Pico PLUS Chemiluminescent Substrate (ThermoFisher Scientific) following the manufacturer’s instructions. Quantification of band signal intensity was performed with the iBright Imaging system (CL1000, ThermoFisher Scientific).

## ACKNOWLEDGEMENTS

We thank Leo Kerner for his assistance in planthopper maintenance and virus transmission experiments. The North Carolina State University, Department of Entomology and Plant Pathology, was part of a team supporting DARPA’s Insect Allies Program. The views expressed are those of the authors and should not be interpreted as representing the official views or policies of the U.S. Government.

## DATA AVAILABILITY STATEMENT

Data supporting the findings of this study are available within the paper and its supplements.

## Supplementary Figures

**Supplementary Figure 1**. Transmission electron micrographs of thin sections of MMV-GFP infected maize showing MMV-GFP virions particles accumulated in the nucleus and cytoplasm of the image. White arrows indicate virions in the cytoplasm. Bars, 2 µm.

**Supplementary Figure 2**. Microscopy images of green fluorescent protein (GFP) expressed from MMV-GFP infections in the *P. maidis* nymphs and adults were microinjected with MMV-GFP purified virions. At 7 and 12 days post injections, the insects were photographed with a fluorescence microscope. Bars, 1000 µm.

## Supplementary Tables

**Table S1. List of primers used in this study**.

**Table S2**. Stable GFP expression by MMV vector in *N. benthamiana*, maize and planthopper following virus passages.

**Table S3**. The predicted intron splicing sites of MMV L gene. Alternative Splice Site Predictor (ASSP): MMV L splice site prediction. Sequence: MMV L; Sequence length 5769 bp, Acceptor site cutoff: 2.2 and Donor site cutoff: 4.5.

