## Supplementary figures and images for "Rescue of the first Alphanucleorhabdovirus entirely from cloned complementary DNA: an efficient vector for systemic expression of foreign genes in maize and insect vectors"

### Supplementary Figure 1

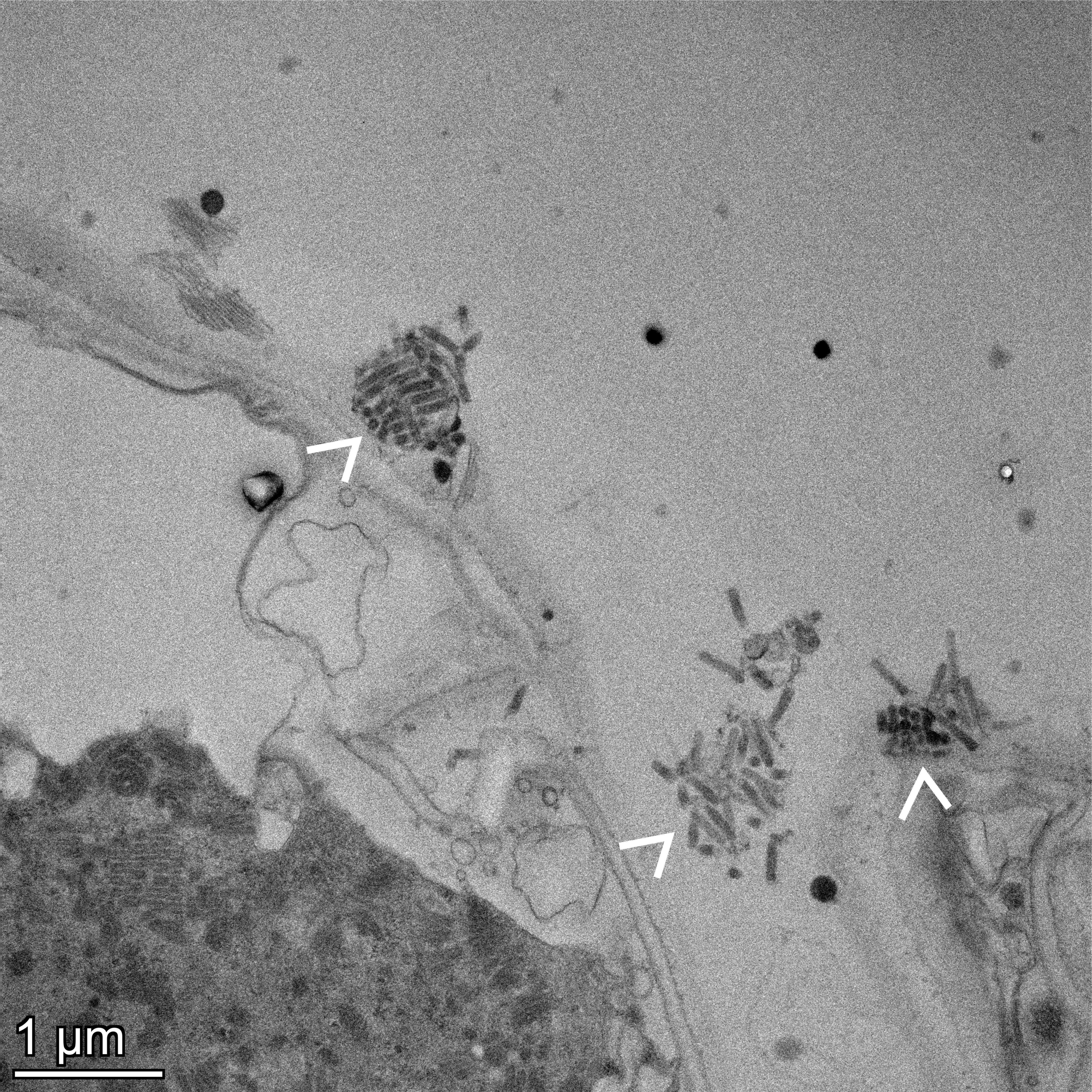

### Supplementary Figure 2

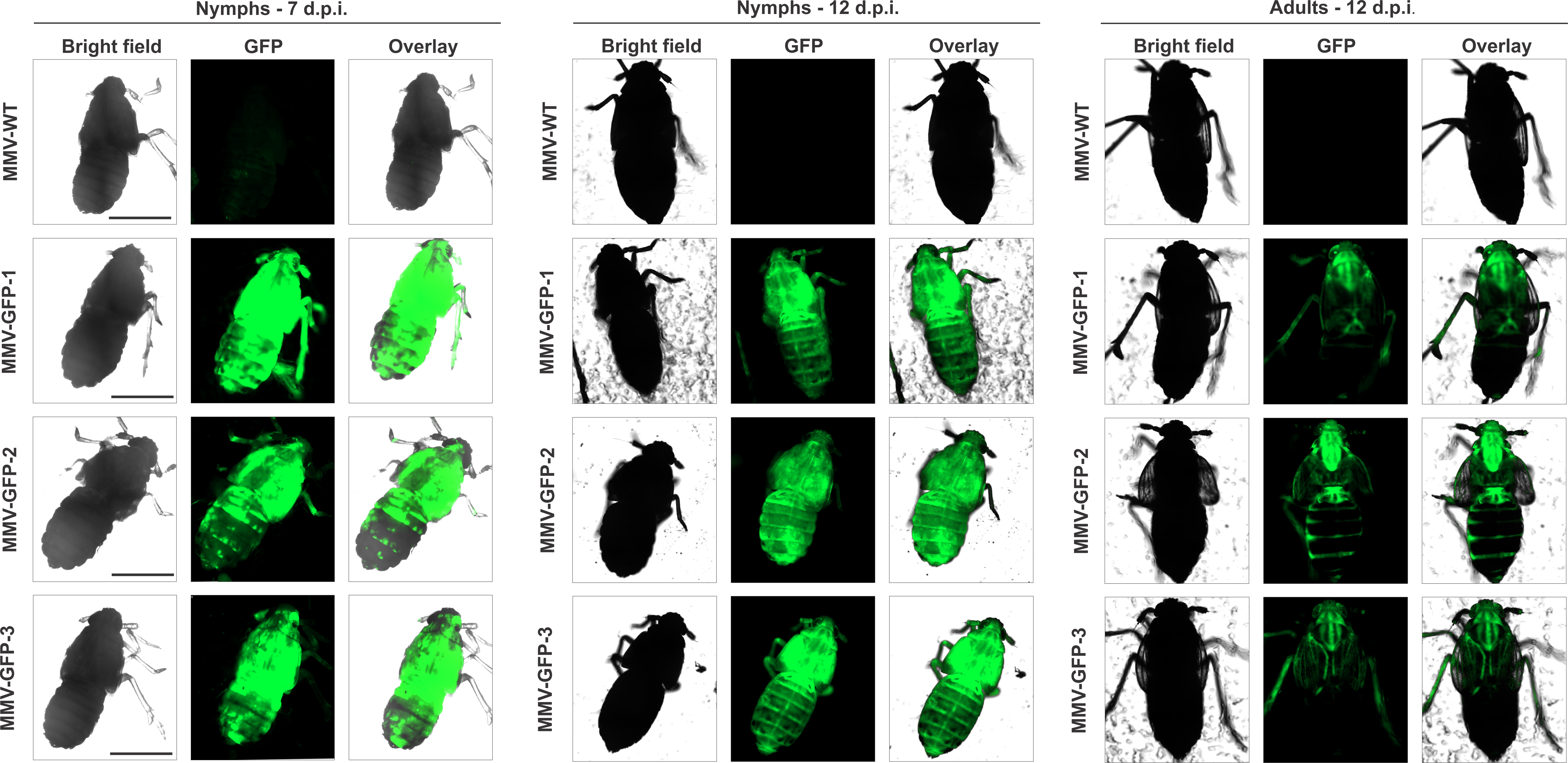
