## Supplementary Table 1 for "Rescue of the first Alphanucleorhabdovirus entirely from cloned complementary DNA: an efficient vector for systemic expression of foreign genes in maize and insect vectors"

**Table S1.** List of primers used in this study.

| **List of primers used for construction of cDNA clones and RT-PCR.** | | |
| --- | --- | --- |
| **Primer** | **Sequence (5' → 3')** | **Application** |
| pJLF | GGGTCGGCATGGCATCTC | Construction of pJL-MMV |
| pJLR | CCTCTCCAAATGAAATGAACTTCC |  |
| pJLMMVF | catttcatttggagaggAGAGACCCATATATTTCATAAA | Construction of pJL-MMV |
| pJLMMVR | gagatgccatgccgacccAGAGACCCAGAAAAACATGGC |  |
| MMVNF | *cacc*ATGGCAAACATCAACATCC | Construction of pTF-N&P and RT-PCR detection of MMV N |
| MMVNR | CTATAAGCCTGATCGTGTCTTC |  |
| MMVPF | *cacc*ATGAATCGTTACTCCCGTCG | Construction of pTF-N&P |
| MMVPR | TTAGATCTCAACCCTAGGGC |  |
| 35SLF | catttcatttggagaggATGGATCCAGATTATCCTGATC | Construction of pJL-L-intron |
| TerLF | AATTCCTAAAACCAAAATCCAGT | Construction of pJL-MMV-intron and pJL-L-intron |
| TerLR | gagatgccatgccgacccTCACCCGAGCAGGTCGGACAC | Construction of pJL-L-intron |
| MMVLXSR | GTCGGATCTCATTCAGTCCATGTGTGGGGGCGTC | Construction of pJL-MMV-intron and pJL-L-intron |
| LXSF | GACGCCCCCACACATGGACTGAATGAGATCCGAC |  |
| MMV1355F | TGTCTGCAGGTGGAGACGAGGCA | Detection of gene inserts between N and P |
| GFP F | ATGGTGAGCAAGGGCGAG | RT-PCR detection of GFP |
| GFP R | CTTGTACAGCTCGTCC |  |
| *Zmactin* F | GGTGGCTCCACTATGTTCCC | RT-PCR detection of maize actin (NCBI accession number: NM_001154731) |
| *Zmactin* R | TATACTGCTCCAGAAATTCA |  |
| *PmeIF1* F | GACGGTTTAGTTCACATAAG | RT-PCR detection of *P. maidis* elongation Factor 1 |
| *PmeIF1* R | TGAACTTTGAGCTGGTC |  |
| **Primers selected for MMV and GFP gene expression analysis.** | | |
| **Primer** | **Sequence (5' → 3')** | **Efficiency (%)** |
| qMMV-N1-Forward | GAAGGCGATCTCATTCCTATG | 99.2 |
| qMMV-N1-Reverse | CCTCTGTCGTATGCTGTTG |  |
| qMMV-N2-Forward | AGCCAACTTTCTCGCATC | 99.9 |
| qMMV-N2-Reverse | GAACCTGCATATCCCATACTC |  |
| qGFP-8-Forward | GCAGAAGAACGGCATCAA | 93.4 |
| qGFP-8-Reverse | TGCTCAGGTAGTGGTTGT |  |
| qGFP-9-Forward | AAGATCCGCCACAACATC | 96.5 |
| qGFP-9-Reverse | ACTGGGTGCTCAGGTAG |  |
| qMMV-P1-Forward | ACTTCACACGACCTTTGC | 95.1 |
| qMMV-P1-Reverse | GTCCTCCATTCCAGTCTTATTC |  |
| qMMV-P2-Forward | GATCCTTGCTGTAGGAATCAG | 97.4 |
| qMMV-P2-Reverse | GAGAGCCACCATTTGAGAC |  |
| qMMV-3_1-Forward | GAGTCCCTGTTCTCAATCTATG | 93.1 |
| qMMV-3_1-Reverse | CCTCGAATCCCGATATTCTTC |  |
| qMMV-3_2-Forward | GGAAGTCGCTTTGTCCTATC | 99.17 |
| qMMV-3_2-Reverse | GGACTCCATAGATTGAGAACAG |  |
| qMMV-M1-Forward | TCTTCGATCCAGACACCTAC | 93.2 |
| qMMV-M1-Reverse | GATTTCCAGACCGTCATCTTC |  |
| qMMV-M7-Forward | TGCTCAGGAAGTATGAGAACC | 96.2 |
| qMMV-M7-Reverse | CCGATGTGTACCCTAGACATATAC |  |
| qMMV-G2-Forward | TCTGGCTGCTACTTCAAATC | 100.1 |
| qMMV-G2-Reverse | GAGCTGAGAAGGCTATATTGAG |  |
| qMMV-G4-Forward | CCCACTCGATTTCGACTATTAC | 97.4 |
| qMMV-G4-Reverse | CGCTTTCTTCTAGTGTGTACC |  |
| qMMV-L3-Forward | GGGTCGGAGATCAACAATAAG | 96.3 |
| qMMV-L3-Reverse | CAGGATCTGGTTTGTCTGTG |  |
| qMMV-L5-Forward | GGTTCCTACACCAAGGAAAG | 94.5 |
| qMMV-L5-Reverse | GCTGCTCCCTTCATCATATC |  |
| qMEP^1^-Forward | TGTACTCGGCAATGCTCTTG | 93.9 |
| qMEP^1^-Reverse | TTTGATGCTCCAGGCTTACC |  |

**Note:** Sequences in lowercase letters are designed to facilitate Hi-fi assembly. Sequences in lowercase and italics are designed to facilitate directional cloning of PCR products into an entry vector for the Gateway system. ^1^ Internal reference gene for maize from Manoli et al., 2012.
