## Supplementary Table 2 for "Rescue of the first Alphanucleorhabdovirus entirely from cloned complementary DNA: an efficient vector for systemic expression of foreign genes in maize and insect vectors"

**Table S2.** Stable GFP expression by MMV vector in *N. benthamiana*, maize and planthopper following virus passages.

| **Experiment** | ***N. benthamiana* to maize** | | | **maize to maize** | | | **maize to *P. maidis*** | | ***P. maidis* to maize** | | |
| --- | --- | --- | --- | --- | --- | --- | --- | --- | --- | --- | --- |
|  | ELISA MMV-GFP Infected plants/inoculated^a^ | RT-PCR MMV-N positive/tested^b^ | RT-PCR GFP positive/tested^c^ | ELISA MMV-GFP Infected/inoculated^d^ | RT-PCR MMV-N positive/tested^b^ | RT-PCR GFP positive/tested^c^ | RT-PCR MMV-N positive/tested^e^ | RT-PCR GFP positive/tested^f^ | ELISA MMV-GFP Infected plants/inoculated^g^ | RT-PCR MMV-N positive/tested^b^ | RT-PCR GFP positive/tested^c^ |
| 1 | 4/80 | 4/4 | 4/4 | 6/120 | 6/6 | 6/6 | 3/10 | 3/10 | 12/20 (60%) | 12/12 | 12/12 |
| 2 | 6/80 | 6/6 | 6/6 | 8/100 | 8/8 | 8/8 | 4/10 | 4/10 | 13/20 (65%) | 13/13 | 13/13 |
| 3 | 4/80 | 4/4 | 4/4 | 9/120 | 9/9 | 9/9 | 4/10 | 4/10 | 14/20 (70%) | 14/14 | 14/14 |

Note: ^a^No. of maize plants infected via VPI/No. of plants VPI with *N. benthamiana* crude extract harbouring MMV-GFP+NPL; ^b^No. of symptomatic maize samples positive for MMV/No. of symptomatic plants tested; ^c^No. of symptomatic maize samples positive for GFP/No .of symptomatic plants tested, only symptomatic plants were tested; ^d^No. of maize plants infected via VPI/No. of plants VPI with maize crude extract harbouring MMV-GFP+NPL; ^e^No. of planthoppers tested positive for virus via MMV-GFP crude extract injections/No. of planthoppers tested by RT-PCR; ^f^No. of planthoppers tested positive for GFP via MMV-GFP crude extract injections/No. of planthoppers tested by RT-PCR; ^g^No. of maize plants infected via insect transmission. Plants tested virus positive/No. of plants tested (Percentage of MMV-GFP infection). Virus infection of the experimental plants were determined by DAS-ELISA using commercial antibodies against MMV (Agdia, Inc. Elkhart, IN, USA) as per the manufacturer's guidelines. For all experiments, only ELISA positive plants were subjected to RT-PCR analysis. MMV N and GFP primers (Table S1) to detect the virus presence and stability of the GFP insertion.
