## Supplementary Table 3 for "Rescue of the first Alphanucleorhabdovirus entirely from cloned complementary DNA: an efficient vector for systemic expression of foreign genes in maize and insect vectors"

**Table S3.** The predicted intron splicing sites of MMV L gene. Alternative Splice Site Predictor (ASSP): MMV L splice site prediction. Sequence: MMV L; Sequence length 5769 bp, Acceptor site cutoff: 2.2 and Donor site cutoff: 4.5.

|  |  |  |  |  | **Activations**** | |  |
| --- | --- | --- | --- | --- | --- | --- | --- |
| **Position (bp)** | **Putative splice site** | **Sequence** | **Score*** | **Intron GC*** | **Alt./Cryptic** | **Constitutive** | **Confidence**** |
| 65 | Alt. isoform/cryptic donor | TACAAGAAGGgtacgatgag | 7.479 | 0.514 | 0.737 | 0.200 | 0.729 |
| 131 | Alt. isoform/cryptic donor | ACCACCTTAAgtcagcactc | 5.062 | 0.486 | 0.939 | 0.043 | 0.954 |
| 155 | Alt. isoform/cryptic acceptor | tcgtacacagATATGATGAA | 2.632 | 0.543 | 0.645 | 0.335 | 0.480 |
| 211 | Alt. isoform/cryptic donor | ATTAACTTTGgtatatctca | 5.192 | 0.414 | 0.929 | 0.049 | 0.947 |
| 241 | Alt. isoform/cryptic acceptor | gatgtcccagATAGAAACAA | 3.570 | 0.429 | 0.593 | 0.384 | 0.352 |
| 268 | Alt. isoform/cryptic acceptor | catcctgaagAGTACTCCAT | 4.171 | 0.386 | 0.879 | 0.116 | 0.868 |
| 304 | Alt. isoform/cryptic donor | ACCCTTTATGgtgatctctt | 5.670 | 0.486 | 0.918 | 0.059 | 0.936 |
| 325 | Alt. isoform/cryptic acceptor | cactcgtcagAGCACGCTCG | 7.210 | 0.471 | 0.850 | 0.141 | 0.835 |
| 404 | Constitutive acceptor | ctcctcctagAGATGACACC | 7.478 | 0.471 | 0.168 | 0.823 | 0.796 |
| 450 | Alt. isoform/cryptic acceptor | ttgctgacagGAAGACATAT | 5.368 | 0.514 | 0.506 | 0.480 | 0.052 |
| 530 | Alt. isoform/cryptic acceptor | gcttggacagAGAAGGGGGC | 2.279 | 0.457 | 0.958 | 0.038 | 0.960 |
| 559 | Alt. isoform/cryptic donor | GATTGGCCTAgtgtgtctct | 7.186 | 0.443 | 0.914 | 0.061 | 0.934 |
| 572 | Alt. isoform/cryptic acceptor | gtgtctctagACAAGAAGAG | 4.334 | 0.457 | 0.563 | 0.421 | 0.252 |
| 580 | Alt. isoform/cryptic donor | GACAAGAAGAgtgggtacgt | 7.413 | 0.414 | 0.929 | 0.051 | 0.945 |
| 584 | Alt. isoform/cryptic donor | AGAAGAGTGGgtacgtgaaa | 7.392 | 0.400 | 0.866 | 0.097 | 0.888 |
| 604 | Alt. isoform/cryptic donor | GTGATGGCAGgtaaagatac | 10.680 | 0.371 | 0.794 | 0.146 | 0.816 |
| 708 | Alt. isoform/cryptic donor | AGCTGACAAGgtggcagaaa | 6.463 | 0.400 | 0.835 | 0.123 | 0.853 |
| 726 | Alt. isoform/cryptic donor | AAGGCTGAATgtgcgttact | 8.709 | 0.371 | 0.794 | 0.156 | 0.804 |
| 741 | Alt. isoform/cryptic acceptor | gttactatagTTGTCTTTGT | 3.700 | 0.457 | 0.776 | 0.213 | 0.725 |
| 757 | Alt. isoform/cryptic acceptor | ttgtgatcagCTTGATATAC | 5.030 | 0.429 | 0.786 | 0.206 | 0.738 |
| 770 | Alt. isoform/cryptic acceptor | gatataccagATAGGATAAC | 2.427 | 0.443 | 0.835 | 0.159 | 0.809 |
| 807 | Alt. isoform/cryptic donor | AATCATCAAAgtgggggatg | 5.996 | 0.471 | 0.768 | 0.181 | 0.765 |
| 852 | Alt. isoform/cryptic donor | CGGATACAATgtaattggat | 6.294 | 0.471 | 0.713 | 0.213 | 0.701 |
| 884 | Constitutive acceptor | ttgcttgcagGAATAATCCA | 7.339 | 0.443 | 0.342 | 0.634 | 0.460 |
| 978 | Alt. isoform/cryptic acceptor | gccccggcagGAAGTATCTA | 3.600 | 0.471 | 0.759 | 0.233 | 0.694 |
| 980 | Alt. isoform/cryptic donor | CCGGCAGGAAgtatctagaa | 5.454 | 0.443 | 0.749 | 0.189 | 0.747 |
| 1110 | Alt. isoform/cryptic donor | GATGCAAGAGgtggcatctg | 7.503 | 0.400 | 0.757 | 0.182 | 0.760 |
| 1200 | Alt. isoform/cryptic acceptor | catatcatagGATACACAAG | 5.484 | 0.343 | 0.579 | 0.405 | 0.301 |
| 1273 | Alt. isoform/cryptic acceptor | ctgtctaaagGATAACTTGA | 3.148 | 0.386 | 0.609 | 0.370 | 0.392 |
| 1318 | Constitutive acceptor | taacttccagGATTGGGACT | 7.849 | 0.371 | 0.358 | 0.620 | 0.423 |
| 1353 | Alt. isoform/cryptic donor | AAACTTCGAAgtgcctttct | 6.165 | 0.429 | 0.904 | 0.070 | 0.923 |
| 1413 | Alt. isoform/cryptic acceptor | ccccgaacagGCAGGAGATT | 6.420 | 0.400 | 0.616 | 0.366 | 0.405 |
| 1444 | Alt. isoform/cryptic donor | TCAACTCGTGgtacaatatt | 7.155 | 0.500 | 0.786 | 0.161 | 0.796 |
| 1465 | Alt. isoform/cryptic acceptor | cgggtctcagAACCGCAGAG | 3.518 | 0.471 | 0.894 | 0.102 | 0.886 |
| 1529 | Alt. isoform/cryptic acceptor | tttcttcaagGCATAAATGA | 2.963 | 0.486 | 0.580 | 0.400 | 0.311 |
| 1588 | Alt. isoform/cryptic donor | CCTAAAGAGCgtgagctgaa | 4.902 | 0.471 | 0.868 | 0.100 | 0.884 |
| 1641 | Alt. isoform/cryptic acceptor | tccggctcagATTGTACTGT | 4.397 | 0.500 | 0.531 | 0.456 | 0.141 |
| 1648 | Alt. isoform/cryptic donor | AGATTGTACTgtgtgtcaac | 6.195 | 0.400 | 0.785 | 0.163 | 0.792 |
| 1661 | Alt. isoform/cryptic acceptor | gtgtcaacagAGGCATTATT | 3.138 | 0.471 | 0.784 | 0.209 | 0.733 |
| 1688 | Alt. isoform/cryptic donor | AAATACTCAAgtatttcccg | 6.768 | 0.400 | 0.895 | 0.077 | 0.914 |
| 1749 | Alt. isoform/cryptic donor | GATGTTCAAAgtgtcctccc | 6.511 | 0.400 | 0.790 | 0.157 | 0.802 |
| 1834 | Alt. isoform/cryptic donor | CAGATGAGACgtaatatatg | 4.604 | 0.386 | 0.864 | 0.096 | 0.889 |
| 1843 | Alt. isoform/cryptic donor | CGTAATATATgtgagcctgt | 8.428 | 0.429 | 0.828 | 0.122 | 0.852 |
| 1867 | Alt. isoform/cryptic donor | TCACAATTAGgtaaactctt | 10.182 | 0.429 | 0.510 | 0.391 | 0.233 |
| 1868 | Constitutive acceptor | tcacaattagGTAAACTCTT | 6.092 | 0.371 | 0.456 | 0.529 | 0.137 |
| 1902 | Constitutive acceptor | tgttcaatagGACTCATGAG | 2.679 | 0.386 | 0.469 | 0.509 | 0.079 |
| 2106 | Alt. isoform/cryptic donor | CAATGTTGATgtatcactaa | 4.523 | 0.457 | 0.917 | 0.057 | 0.937 |
| 2143 | Alt. isoform/cryptic donor | CAAGTCCTTGgtatcaccat | 5.238 | 0.514 | 0.899 | 0.070 | 0.922 |
| 2159 | Alt. isoform/cryptic acceptor | accatatcagGAGTGACCAG | 3.107 | 0.429 | 0.669 | 0.316 | 0.528 |
| 2217 | Alt. isoform/cryptic acceptor | tcgcccgaagGACAATCAAG | 4.142 | 0.514 | 0.682 | 0.298 | 0.564 |
| 2295 | Alt. isoform/cryptic donor | TGAAACATGGgtcagtgact | 8.950 | 0.414 | 0.842 | 0.119 | 0.859 |
| 2315 | Alt. isoform/cryptic donor | CTCTCTTCATgtacaacaaa | 4.522 | 0.414 | 0.945 | 0.039 | 0.959 |
| 2355 | Alt. isoform/cryptic acceptor | ttcctctcagATCACCTCTG | 3.901 | 0.414 | 0.616 | 0.371 | 0.398 |
| 2565 | unclassified acceptor | cttgcttcagGACACCAACA | 3.664 | 0.471 | 0.478 | 0.490 | 0.000 |
| 2625 | Alt. isoform/cryptic donor | ATCAGTGAAGgtgactagga | 8.621 | 0.443 | 0.523 | 0.385 | 0.264 |
| 2667 | Alt. isoform/cryptic donor | GTTATGTATGgtgggatctt | 7.844 | 0.486 | 0.812 | 0.140 | 0.828 |
| 2695 | Alt. isoform/cryptic donor | GGACTACCTGgtactataca | 6.690 | 0.443 | 0.852 | 0.110 | 0.871 |
| 2771 | Alt. isoform/cryptic acceptor | ttcatttcagAATTGGTTCC | 7.486 | 0.443 | 0.596 | 0.390 | 0.345 |
| 2852 | Alt. isoform/cryptic donor | GAATGACAGAgtatgcgaaa | 5.216 | 0.471 | 0.861 | 0.102 | 0.882 |
| 3002 | Constitutive acceptor | accctcctagACAGACAGAG | 8.560 | 0.514 | 0.407 | 0.577 | 0.295 |
| 3006 | Alt. isoform/cryptic acceptor | tcctagacagACAGAGAGAA | 3.489 | 0.514 | 0.655 | 0.327 | 0.501 |
| 3063 | Alt. isoform/cryptic donor | GGATGTGAAGgtgctgcacg | 6.990 | 0.500 | 0.917 | 0.062 | 0.933 |
| 3075 | Alt. isoform/cryptic donor | GCTGCACGAAgtagccggag | 5.352 | 0.486 | 0.937 | 0.044 | 0.953 |
| 3133 | Alt. isoform/cryptic acceptor | tattgaccagACAAGAACGG | 3.696 | 0.514 | 0.884 | 0.110 | 0.875 |
| 3141 | Alt. isoform/cryptic donor | GACAAGAACGgtgagagcgt | 9.114 | 0.429 | 0.754 | 0.193 | 0.744 |
| 3225 | Alt. isoform/cryptic acceptor | tgttcgctagGGACATACAT | 4.476 | 0.429 | 0.561 | 0.422 | 0.247 |
| **3281** | **Alt. isoform/cryptic donor** | **CTGCGGACAGgtatcgatat** | **8.521** | **0.400** | **0.773** | **0.164** | **0.788** |
| 3318 | Alt. isoform/cryptic donor | GGTTTTAGGGgtaacagtac | 7.729 | 0.514 | 0.950 | 0.034 | 0.964 |
| 3460 | Alt. isoform/cryptic donor | ATATATCAAGgttcctacac | 5.704 | 0.543 | 0.957 | 0.031 | 0.968 |
| 3475 | Alt. isoform/cryptic acceptor | ctacaccaagGAAAGGTTCA | 2.238 | 0.443 | 0.871 | 0.124 | 0.858 |
| 3479 | Alt. isoform/cryptic donor | CCAAGGAAAGgttcaaggcc | 5.027 | 0.557 | 0.951 | 0.034 | 0.964 |
| 3615 | Alt. isoform/cryptic acceptor | ttgctctcagAGCCATCACT | 5.802 | 0.414 | 0.633 | 0.351 | 0.446 |
| 3789 | Alt. isoform/cryptic acceptor | cacatagcagGGGAGGGAAA | 3.130 | 0.471 | 0.666 | 0.320 | 0.519 |
| 3820 | Alt. isoform/cryptic acceptor | ccatttccagAGTGTACTGA | 5.296 | 0.457 | 0.628 | 0.348 | 0.446 |
| 3846 | Alt. isoform/cryptic acceptor | tattatacagAGCCATGGTC | 4.104 | 0.400 | 0.780 | 0.204 | 0.738 |
| 3884 | Alt. isoform/cryptic acceptor | ggtctaccagAGATGGTCTG | 4.183 | 0.471 | 0.797 | 0.196 | 0.754 |
| 3893 | Alt. isoform/cryptic donor | AGATGGTCTGgtattcaaaa | 5.523 | 0.429 | 0.841 | 0.114 | 0.864 |
| 3950 | Alt. isoform/cryptic acceptor | ccgtctccagAACGAACAAC | 2.959 | 0.443 | 0.741 | 0.246 | 0.668 |
| 3974 | Alt. isoform/cryptic acceptor | gctttgtcagAGATGCCATC | 4.202 | 0.471 | 0.753 | 0.235 | 0.687 |
| 4013 | Alt. isoform/cryptic acceptor | gcattcatagAGTCAGTAAA | 7.565 | 0.457 | 0.613 | 0.375 | 0.389 |
| 4013 | Alt. isoform/cryptic donor | CATTCATAGAgtcagtaaat | 6.676 | 0.386 | 0.872 | 0.094 | 0.892 |
| 4023 | Alt. isoform/cryptic donor | GTCAGTAAATgtgaagttgg | 4.781 | 0.386 | 0.937 | 0.044 | 0.953 |
| 4047 | Alt. isoform/cryptic donor | TCATCACCAGgttgagatag | 6.340 | 0.400 | 0.836 | 0.121 | 0.855 |
| 4048 | Alt. isoform/cryptic acceptor | tcatcaccagGTTGAGATAG | 4.956 | 0.400 | 0.692 | 0.295 | 0.574 |
| 4117 | Alt. isoform/cryptic donor | TCTTTAACAAgtgtggatga | 6.384 | 0.471 | 0.918 | 0.058 | 0.937 |
| 4170 | Alt. isoform/cryptic donor | ATCCCGTCAAgtgtcagaat | 4.713 | 0.443 | 0.860 | 0.107 | 0.875 |
| 4260 | Alt. isoform/cryptic acceptor | tccttgtcagGAAACTCGGA | 5.201 | 0.471 | 0.772 | 0.221 | 0.714 |
| 4293 | Alt. isoform/cryptic acceptor | tccctataagAAACACCAAA | 3.820 | 0.443 | 0.726 | 0.259 | 0.644 |
| 4476 | Alt. isoform/cryptic donor | TCTTCAAAATgtgcctctcc | 6.853 | 0.529 | 0.840 | 0.119 | 0.858 |
| 4489 | Alt. isoform/cryptic acceptor | gcctctccagCTGGCCTTGG | 5.712 | 0.400 | 0.607 | 0.381 | 0.373 |
| 4525 | Alt. isoform/cryptic donor | CCTGATGTTTgtgagtgtct | 10.324 | 0.443 | 0.634 | 0.292 | 0.540 |
| 4545 | Alt. isoform/cryptic acceptor | ttgagtgcagAGCCATCATT | 3.307 | 0.514 | 0.950 | 0.048 | 0.949 |
| 4578 | Alt. isoform/cryptic donor | TAAGAGGATGgtagggcaat | 7.778 | 0.500 | 0.770 | 0.166 | 0.784 |
| 4654 | Alt. isoform/cryptic donor | AGGGTCCACAgtgagaggat | 7.162 | 0.429 | 0.639 | 0.286 | 0.553 |
| 4822 | Constitutive acceptor | cctgcctcagCTGTATATTA | 7.563 | 0.557 | 0.362 | 0.613 | 0.410 |
| 4848 | Alt. isoform/cryptic donor | GGCCTGCCTGgtttgtcttc | 6.531 | 0.500 | 0.897 | 0.077 | 0.914 |
| 4867 | Alt. isoform/cryptic acceptor | tcccttggagGGTGTGATTG | 4.025 | 0.486 | 0.514 | 0.464 | 0.096 |
| 4867 | Alt. isoform/cryptic donor | CCCTTGGAGGgtgtgattgt | 6.631 | 0.514 | 0.932 | 0.048 | 0.948 |
| 4905 | Alt. isoform/cryptic donor | GATCAATCTGgtgtccaagg | 5.752 | 0.557 | 0.757 | 0.190 | 0.749 |
| 4914 | Alt. isoform/cryptic donor | GGTGTCCAAGgtgttggtcg | 6.723 | 0.557 | 0.787 | 0.162 | 0.794 |
| 4944 | Alt. isoform/cryptic donor | GACCATCCAGgtctatatca | 5.263 | 0.486 | 0.887 | 0.081 | 0.908 |
| 5018 | Alt. isoform/cryptic acceptor | ctgcttccagAGGATCTTAG | 4.973 | 0.486 | 0.847 | 0.147 | 0.827 |
| 5057 | Alt. isoform/cryptic acceptor | acaaccacagACAAACCAGA | 2.736 | 0.443 | 0.698 | 0.286 | 0.591 |
| 5093 | Alt. isoform/cryptic acceptor | caactggcagCGGATGACAC | 2.317 | 0.500 | 0.869 | 0.122 | 0.860 |
| 5166 | Alt. isoform/cryptic donor | GACGAGGAAGgtgcacggaa | 4.630 | 0.514 | 0.909 | 0.067 | 0.926 |
| 5219 | Alt. isoform/cryptic acceptor | ggattcgcagGGGTCATGGT | 3.265 | 0.529 | 0.743 | 0.247 | 0.668 |
| 5226 | Alt. isoform/cryptic donor | AGGGGTCATGgtgacagatc | 6.523 | 0.500 | 0.837 | 0.122 | 0.854 |
| 5360 | Alt. isoform/cryptic acceptor | gtgttggcagGTTCTAGACT | 4.759 | 0.414 | 0.745 | 0.239 | 0.679 |
| 5414 | Alt. isoform/cryptic acceptor | tcgcctgcagGCCTAGAGAT | 3.848 | 0.586 | 0.856 | 0.134 | 0.844 |
| 5436 | Alt. isoform/cryptic donor | TGTCAGGAAAgtgagggcaa | 6.941 | 0.543 | 0.895 | 0.077 | 0.914 |
| 5534 | Alt. isoform/cryptic donor | TGATCTCCCTgtatgctcat | 5.251 | 0.529 | 0.905 | 0.070 | 0.922 |
| 5586 | Alt. isoform/cryptic donor | CCTTGTCAAGgtgaacatac | 8.154 | 0.600 | 0.760 | 0.182 | 0.760 |
| 5622 | Alt. isoform/cryptic donor | TGTCACCGCTgtgtgcgacc | 6.619 | 0.586 | 0.807 | 0.152 | 0.811 |
| 5681 | Alt. isoform/cryptic acceptor | ttcctaaaagAAAAGGCTCC | 4.668 | 0.600 | 0.898 | 0.098 | 0.891 |
| 5712 | Alt. isoform/cryptic acceptor | tcattctaagGAGCAGGTTG | 6.368 | 0.486 | 0.725 | 0.261 | 0.640 |

* Scores of the preprocessing models reflecting splice site strength, i.e. a PSSM for putative acceptor sites, and an MDD model for putative donor sites. Intron GC values correspond to 70 nt of the neighboring intron.

** Activations are output values of the backpropagation networks used for classification. High values for one class with low values of the other class imply a good classification. Confidence is a simple measure expressing the differences between output activations. Confidence ranges between zero (undecided) to one (perfect classification).
